# Cicada endosymbionts contain tRNAs that are processed despite having genomes that lack tRNA processing activities

**DOI:** 10.1101/365791

**Authors:** James T. Van Leuven, Meng Mao, Gordon M. Bennett, John P. McCutcheon

## Abstract

Gene loss and genome reduction are defining characteristics of nutritional endosymbiotic bacteria. In extreme cases, even essential genes related to core cellular processes such as replication, transcription, and translation are lost from endosymbiont genomes. Computational predictions on the genomes of the two bacterial symbionts of the cicada *Diceroprocta semicincta, “Candidatus* Hodgkinia cicadicola” (*Alphaproteobacteria*) and “*Ca*. Sulcia muelleri” (*Betaproteobacteria*), find only 26 and 16 tRNA, and 15 and 10 aminoacyl tRNA synthetase genes, respectively. Furthermore, the original “*Ca.* Hodgkinia” genome annotation is missing several essential genes involved in tRNA processing, such as RNase P and CCA tRNA nucleotidyltransferase, as well as several RNA editing enzymes required for tRNA maturation. How “*Ca*. Sulcia” and “*Ca*. Hodgkinia” preform basic translation-related processes without these genes remains unknown. Here, by sequencing eukaryotic mRNA and total small RNA, we show that the limited tRNA set predicted by computational annotation of “*Ca*. Sulcia” and “*Ca*. Hodgkinia” is likely correct. Furthermore, we show that despite the absence of genes encoding tRNA processing activities in the symbiont genomes, symbiont tRNAs have correctly processed 5’ and 3’ ends, and seem to undergo nucleotide modification. Surprisingly, we find that most “*Ca*. Hodgkinia”and “*Ca*. Sulcia” tRNAs exist as tRNA halves. Finally, and in contrast with other related insects, we show that cicadas have experienced little horizontal gene transfer that might complement the activities missing from the endosymbiont genomes. We conclude that “*Ca*. Sulcia” and “*Ca*. Hodgkinia” tRNAs likely function in bacterial translation, but require host-encoded enzymes to do so.

## INTRODUCTION

Bacterial genomes experience structural and genome composition changes during the transition from a free-living to an intracellular lifestyle (1, 2). At the onset of endosymbiosis, symbiont genomes undergo genome rearrangement, mobile element proliferation, and pseudogenization of non-essential genes (3, 4). Following this period of genomic turmoil, endosymbionts evolve towards structural and functional stability while continuing to lose non-coding DNA and genes not critical to symbiont function (5). The resulting small, gene-dense genomes are often stable in gene order and orientation, but experience rapid sequence evolution that is likely caused by the loss of recombination and DNA-repair machinery and sustained reductions in effective population size (1, 6–10). The most gene-poor endosymbiont genomes have lost even seemingly essential genes, like those involved in DNA replication and translation (11). In terms of genome size and coding capacity, these tiny genomes span the gap between their less degenerate endosymbiotic cousins, which retain seemingly minimal sets of genes, and the mitochondria and plastids, which have lost most genes involved in replication, transcription, and translation (11, 12). These extremely gene-poor insect endosymbiont genomes thus provide an opportunity to learn more about key adaptations enabling co-dependent and integrated endosymbioses, but in associations that are younger and more labile than the classic cellular organelles.

*“Candidatus* Hodgkinia cicadacola” (*Alphaproteobacteria*) and *“Ca.* Sulcia muelleri” (*Bacteroidetes;* hereafter *Hodgkinia* and *Sulcia*) have two of the smallest bacterial genomes published. *Sulcia* and *Hodgkinia* are obligate nutritional endosymbionts of many cicadas (13, 14). In the cicada *Diceroprocta semicincta*, the *Hodgkinia* genome is 143 kilobases (kb) and the *Sulcia* genome is 277 kb. Together these reduced endosymbiont genomes encode complementary gene pathways to make the ten essential amino acids required by their cicada host (15). Only 16 tRNA genes and 10 of the 20 required aminoacyl tRNA synthetase genes (aaRSs) were found in the *Hodgkinia* genome using computational methods (16). While the number of tRNA genes encoded in bacterial genomes is quite variable (Fig. 1), theoretical estimates predict that a minimum of ~20 tRNAs are required to translate all 61 codons (17, 18). *Hodgkinia* is missing tRNA genes needed to decode leucine, valine, arginine, serine, threonine, aspartic acid, asparagine, and tyrosine codons (Fig. 2). The mealybug endosymbiont *Candidatus* Tremblaya (hereafter *Tremblaya*) also falls below the theoretical limit of 20, encoding only 8-12 tRNAs genes and 0 or 1 aaRSs, depending on strain (Fig. 1, Table S1) (19–22). However, *Tremblaya* is unusual in hosting its own intrabacterial endosymbiont, which may provide the missing tRNAs and aaRSs (20). There is no such explanation for the apparent lack of tRNA, aaRS, and tRNA processing genes in *Hodgkinia.*

**Figure 1.**
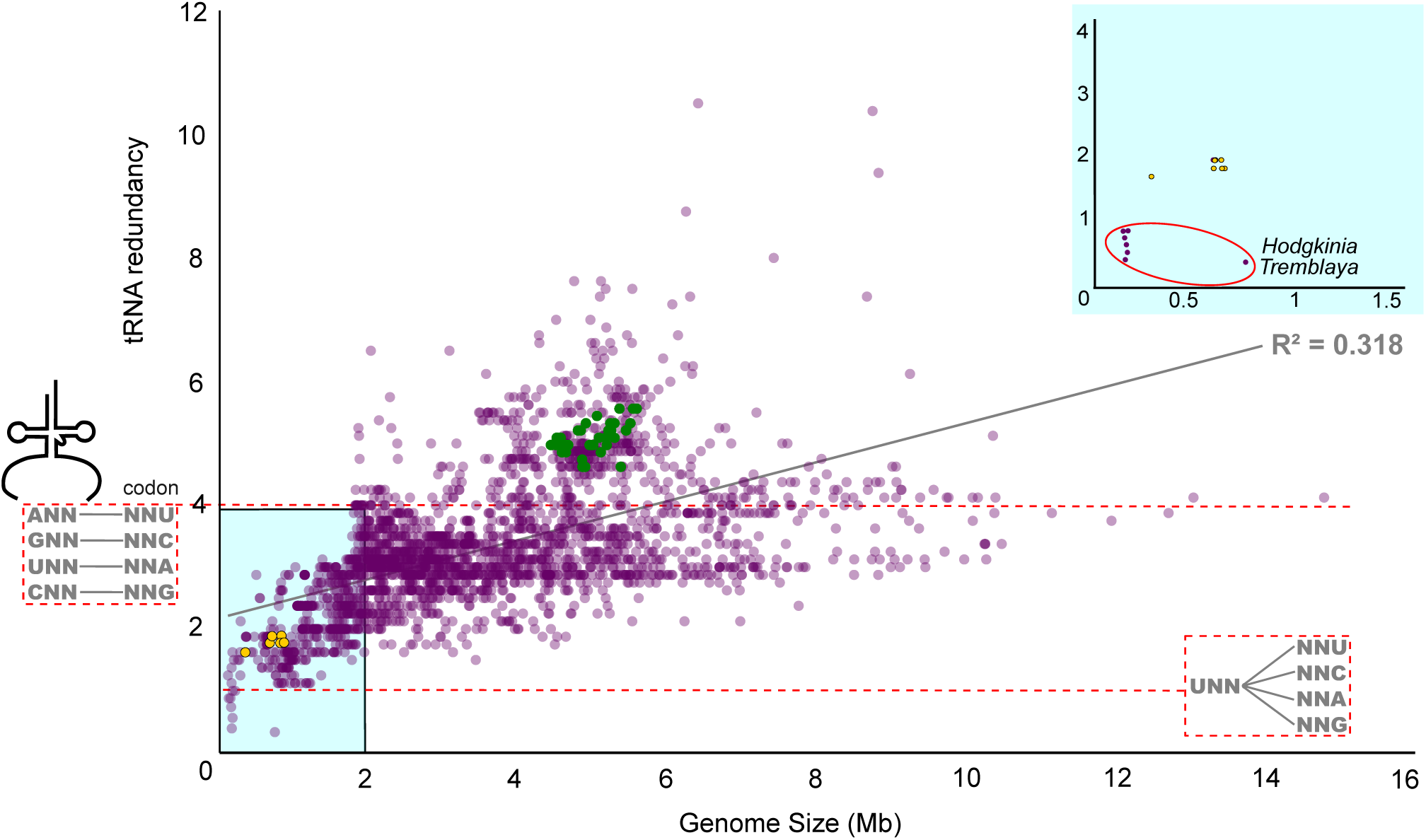
Genome size and tRNA redundancy are positively correlated. Each fully sequenced bacterial genome is shown as a dot (n=2761). tRNA redundancy represents the number of total 4-box tRNA genes in a genome over the number of 4-box families. The red-dashed line at y=1 shows a limit where only one tRNA is found from each of the eight 4-box families. Below this limit, it is unclear if the organism has enough tRNAs for translation. The red-dashed line at y=4 shows one tRNA gene for each 4-box codon. *Buchnera aphidicola* (*Buchnera*) and *Escherichia coli* (*E.coli*) are shown as yellow and green dots, respectively. Theoretically, all bacteria could function with redundancy value of 1.

**Figure 2.**
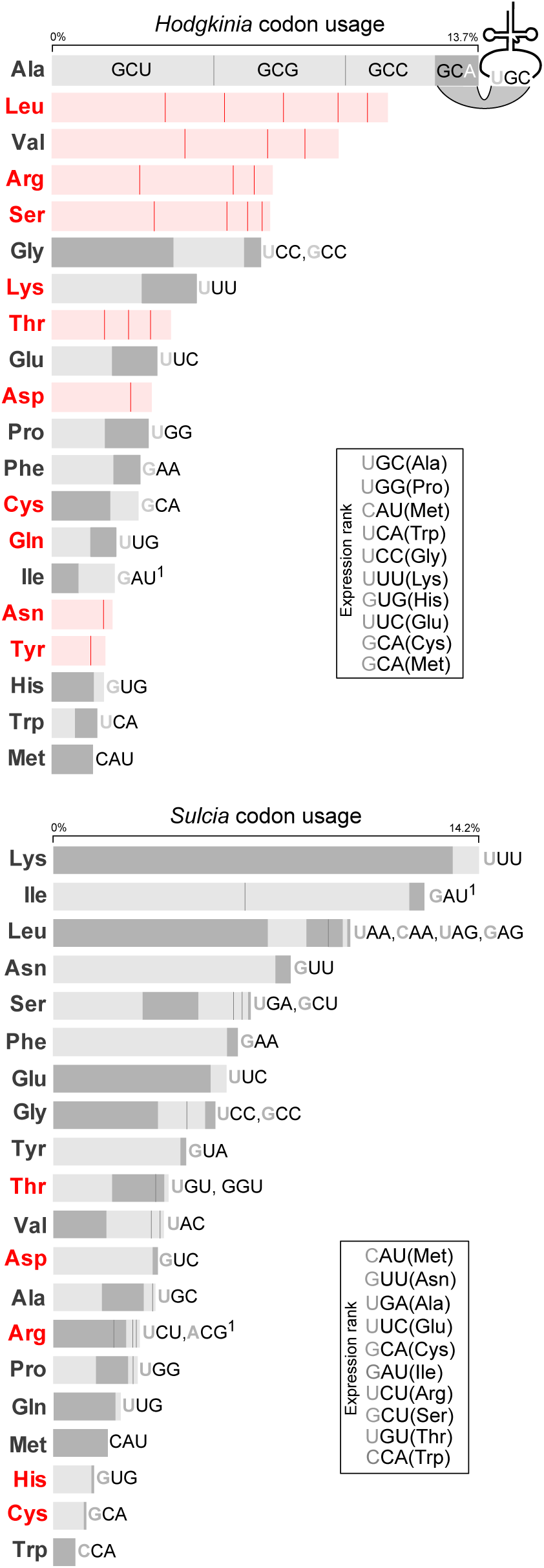
Codon usage and RNA expression in *Hodgkinia* and *Sulcia* genomes. Box size indicates codon frequency of all protein coding genes in *Hodgkinia* and *Sulcia.* Codons are ordered from highest to lowest usage; e.g. of the four alanine codons found in *Hodgkinia* protein coding genes, GCU is used most frequently. The nucleotide sequences for *Hodgkinia* alanine codons (which make up 13.7% of the genome) are shown as an example, all others are omitted for simplicity of display. The presence of a perfectly paired tRNA is indicated by a dark grey box. Light grey fill indicates that a tRNA could possibly be used to translate the codon by N34 wobble. The anticodon sequence of each tRNA is shown to the right of its cognate codons and is written 5’ to 3’. N34 modifications that are likely needed for tRNA-codon pairing are indicated1. A red colored three letter amino acid abbreviation indicates that the genome does not encode that aaRS. tRNA abundance by RNAseq is shown in the “Expression rank” box.

Functional tRNAs are generated by a complex, multistep process that usually requires trimming off transcribed nucleotides that precede (5’ leader) and follow (3’ trailer) the predicted tRNA gene, post-transcriptional nucleotide editing at numerous positions, the addition of a terminal CCA, and aminoacylation of the mature tRNA to produce a molecule that is active on the ribosome. After transcription, 5’ leaders are trimmed by the nearly universal ribozyme RNase P (23, 24). The 3’ trailer is cleaved off by a combination of endonucleases and/or exonucleases (25) and if a terminal 3’ CCA is not encoded in the genome, one is added by a nucleotidyl transferase (26, 27). Finally, tRNA nucleosides are modified by a variety of enzymes at various conserved positions (28–30). These important modifications influence tRNA tertiary structure and interactions with cellular enzymes and proteins (28).

The original published computational annotation of the *Hodgkinia* genome from *D. semicincta* lacks most genes related to tRNA processing (16). It is missing the RNA (*rnpB*) and protein (*rnpA*) subunits of RNase P, and the nucleases responsible for 3’ trailer trimming. *Hodgkinia* does not encode a CCA-adding enzyme, despite having only one tRNA gene with a genome-encoded terminal CCA. This *Hodgkinia* genome contains only three genes involved in tRNA editing (*mnmE*, *mnmA*, and *mnmG*), all of which are predicted to be involved in the conversion of uridine to 5-methylaminomethyl-2-thiouridine at U34 (16, 31). Because the genes encoding aaRSs and tRNA processing enzymes are large and typically highly conserved across life, it is unlikely that these proteins were missed in the original genome annotations.

The dramatic tRNA and aaRS gene loss observed in *Hodgkinia* is extremely rare in bacteria: only *Hodgkinia* and *Tremblaya* lack the theoretical minimum number of tRNAs and aaRS genes in their genomes (Fig. 1, Table S1). The detection of tRNA genes in highly degraded genomes— particularly in mitochondrial genomes—is notoriously difficult (32, 33). Many mitochondrial tRNAs have unusual structures, in some cases missing entire D-loops, making them easy to miss by computational gene finding algorithms unless they are specifically trained to find them (34). For example, the archaeal tRNA genes in the degenerate genome of *Nanoarchaeum* were initially missed because of the unusual way that they are the split and permuted (35, 36). Therefore, we reasoned that our initial computational annotation of *Hodgkinia’s* tRNAs (16) may be incomplete.

Mitochondrial and plastid genomes are missing some to all genes involved in translation. While some organelle genomes are incredibly similar to free-living bacterial genomes (37), mitochondria, and plastids always require extensive genetic complementation by the host (38, 39). Most of the proteins present in organelles are encoded on the host genome, the products of which are imported into the organelle (40). Interestingly, the functioning of some bacterial endosymbionts in amoeba (41–43), protists (44), and insects (20, 45–47) also seem to be supported by horizontal gene transfer (HGT) to the host genomes, although protein import has been established in only a few cases (41, 43, 44, 48). In the insect examples, most transferred genes do not originate from the symbionts themselves, but from other, germ line-associated bacteria (20, 45–47).

The losses of these key genes in *Hodgkinia* raise several questions. Were *Hodgkinia* tRNAs and other small RNAs missed during computational gene prediction? For tRNAs present on the genome, are their 5’ and 3’ ends correctly processed? Are *Hodgkinia* tRNAs modified only at the U34 wobble position as expected based on the gene content of the genome? And, do host enzymes complement the missing symbiont genes? Here we address these questions by sequencing messenger and small RNAs from the cicada species *D. semicincta.*

## RESULTS

### Endosymbiont tRNA genes are correctly annotated

We sequenced small RNAs expressed in cicada bacteriome tissues to experimentally search for unannotated *Hodgkinia* and *Sulcia* tRNAs. Of 145,176,847 quality-filtered reads of length 18-90nts, 15.6% and 28.4% map to the *Hodgkinia* and *Sulcia* genomes, respectively. While the mean genome-wide read coverage of the *Sulcia* and *Hodgkinia* genomes is on the same order (224 reads/bp and 164 reads/bp, respectively), *Hodgkinia* tRNA expression levels are extremely variable (Fig. S1). The comparatively even tRNA, mRNA, and rRNA expression (Fig. 3) from the *Sulcia* genome (compared to *Hodgkinia*) is a feature of unknown significance, but suggests that the near zero expression level for many *Hodgkinia* tRNAs is not due to under sequencing.

**Figure 3.**
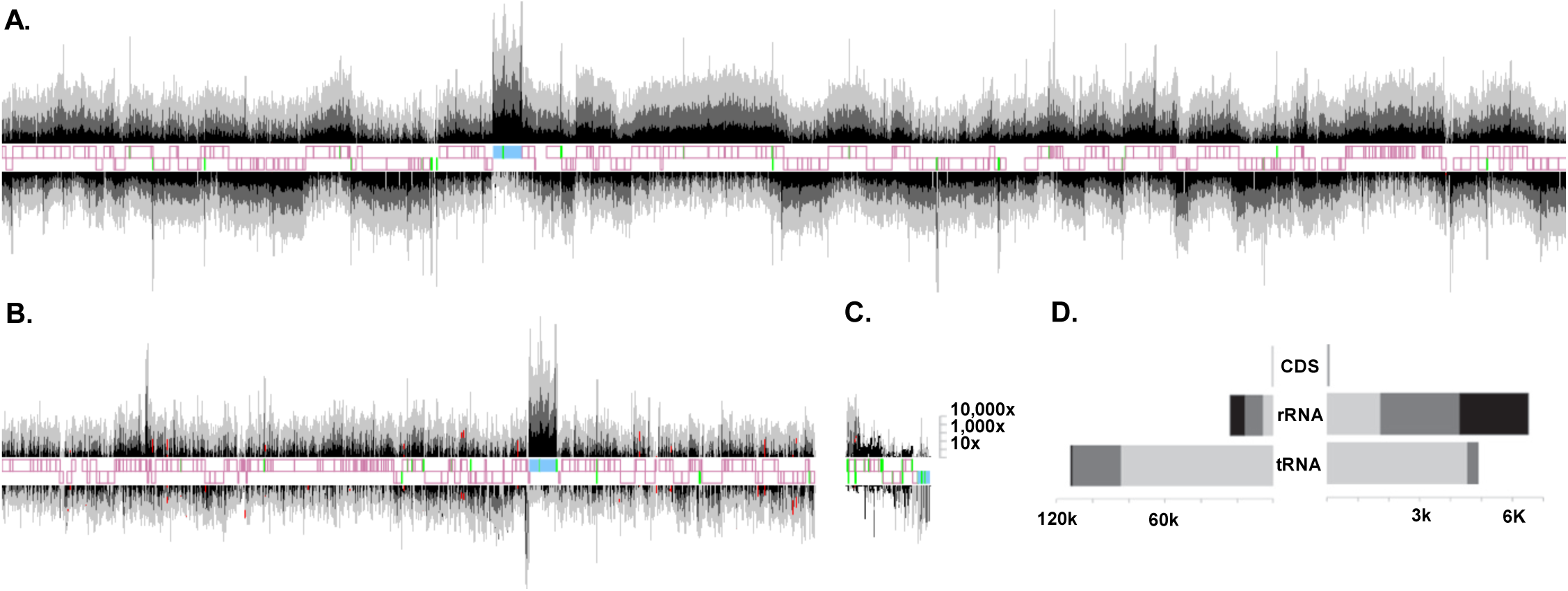
RNA expression patterns from the *Sulcia, Hodgkinia*, and mitochondrial genomes show relatively low expression of *Hodgkinia* tRNAs. Read depth plotted across the *Sulcia* (A), *Hodgkinia* (B) and mitochondrial (C) genomes. Protein coding, ribosomal RNA, and tRNA genes on the sense and anti-sense strands are shown in pink, blue, and green, respectively. Red dots show the highest read depth for each tRNA. Coverage depth for reads of length 18-47, 48-89, and 90-100 are shown in light grey, grey, and black, respectively and each are drawn on a Iog10 scale, then summed. Median coverage depths for *Sulcia* (left) and *Hodgkinia* (right) are shown for each gene category and read length in (D). The bars are colored as in A-C.

In *Sulcia*, we find that >99% of reads map to tRNAs, tmRNA, RNase P, and ribosomal RNAs (Fig. 3, Table S2). The median read depth of reads mapping to the rest of the genome was 380X. To identify unannotated tRNAs, we manually inspected read coverage across the genome to identify regions with pronounced spikes in coverage. Only one of these high coverage spikes were found in intergenic regions (positions 75798-75839). The remaining spikes occur within the bounds of annotated protein-coding sequences (CDSs). The 75,798-75,839 spike corresponds to a Thr^GGT^ tRNA that was unannotated in the original *Sulcia* annotation due to a small error in the published assembly. We updated the *Sulcia* genome to reflect this change (NCBI Reference Sequence NC_012123.1; see Table S3 for primer sequences). In contrast to the tRNA reads, the reads contributing to these CDS spikes do not have terminal CCAs, nor predicted folded structures that resemble tRNA (49). In summary, none of the spikes in coverage can be attributed to unannotated tRNAs.

In *Hodgkinia*, we find high expression of predicted tRNAs, ribosomal RNAs, and the 5’ ends of protein coding genes. Our analysis uncovered previously unannotated RNase P and tmRNA genes (discussed in the “Discovery of unannotated RNase P and tmRNA genes in *Hodgkinia*” section below). Of *Hodgkinia’s* sixteen total tRNA genes, many are not expressed above background (Fig. 4, Table S4). The tRNA genes Gly061 and Gly108 each have no full-length reads aligning to them, even when allowing for 5-8 mismatches to accommodate modified bases (Table S4). However, many shorter-than-full-length reads map to these genes, allowing us to predict modification sites.

**Figure 4.**
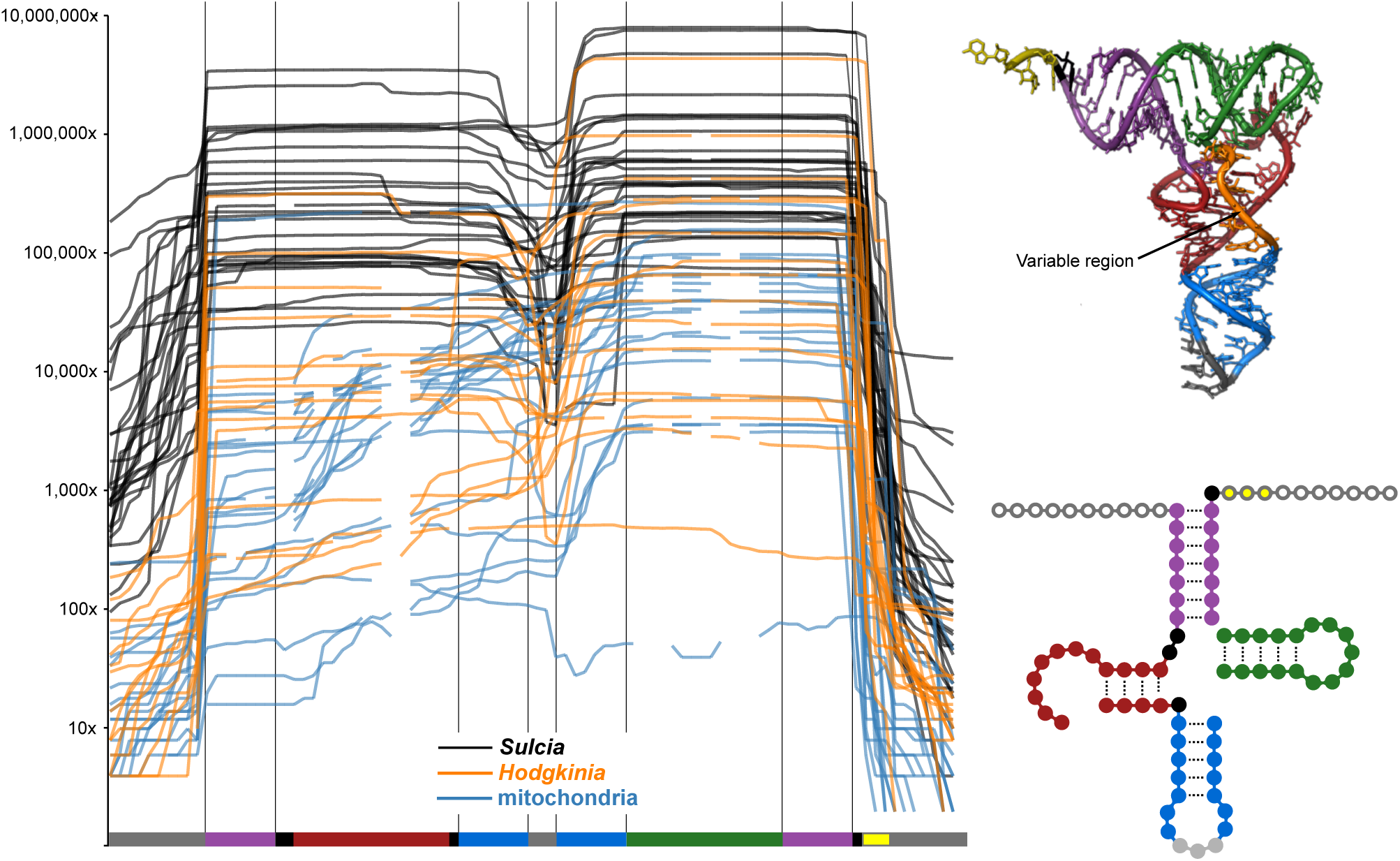
Dynamic range of tRNA half expression in *Sulcia* and *Hodgkinia.* Line graphs show read depth across each tRNA in *Hodgkinia* (orange), *Sulcia* (black), and the cicada mitochondria (blue). Because tRNA length is variable, only bases present in all symbiont tRNA genes are shown in the structural diagrams of tRNAs. Mitochondrial tRNAs are often missing these regions, as indicated by gaps in the line graph. 18-100 nucleotide reads were mapped for this figure.

Mapping reads to endosymbiont genomes allowed us to characterize tRNA processing patterns and uncover expression of unannotated genes, but was not well-suited for identifying spliced or otherwise unconventional RNAs, such as intron containing tRNAs. Therefore, we collapsed identical reads of length 48-90nts and searched for transcripts ending in CCA, containing predicted tRNA genes, or that partially match the *Sulcia* or *Hodgkinia* genomes. Because the number of unique, or nearly unique reads was very high, we choose a minimum coverage cutoff of 100X (see Materials and Methods for justification). The only transcript we found belonging to *Sulcia* or *Hodgkinia* (blast e-value < 1E-25) that was not an annotated tRNA or tmRNA was the previously unannotated *Sulcia* Thr^GGT^ described above. Given these data, we conclude that the computational tRNA predictions for *Hodgkinia* and *Sulcia* are correct, and that these genomes do not encode complete sets of tRNAs.

### Most tRNAs are found as tRNA halves

Of 145,176,847 quality-filtered reads 18-90nts in length, only 0.05% (74,651), and 1.7% (2,520,749) map in full-length to *Hodgkinia* and *Sulcia* tRNAs, respectively (Table S2). Most reads were shorter than the predicted tRNA genes (Figs. 4-6, Table S4). The abundance of short reads could be caused by RNA degradation, PCR bias towards short amplicons during library creation, or bona fide stable tRNA halves (50–53). Because reverse transcription occurs after RNA adapter ligation, these short reads are not likely due to reverse transcriptase failing to proceed through modified nucleotides, because these products would not include both priming sites for PCR. The presence of high levels of tRNA halves was corroborated by cloning and Sanger sequencing a small RNA library prior to PCR amplification. All Sanger sequences align to halves of *Sulcia* tRNA genes and are flanked by adapter and vector sequences (n=9). We conclude that there are large amounts of either tRNA degradation products or stable tRNA halves present in the adult cicada bacteriome. tRNA read coverage is generally highest immediately 3’ of the anticodon and lowest at the anticodon (Fig. 4). Fig. 5 shows that there are exceptions to this pattern, especially in reads mapping to *Sulcia*, where many 5’ halves are more abundant than their 3’ halves.

**Figure 5.**
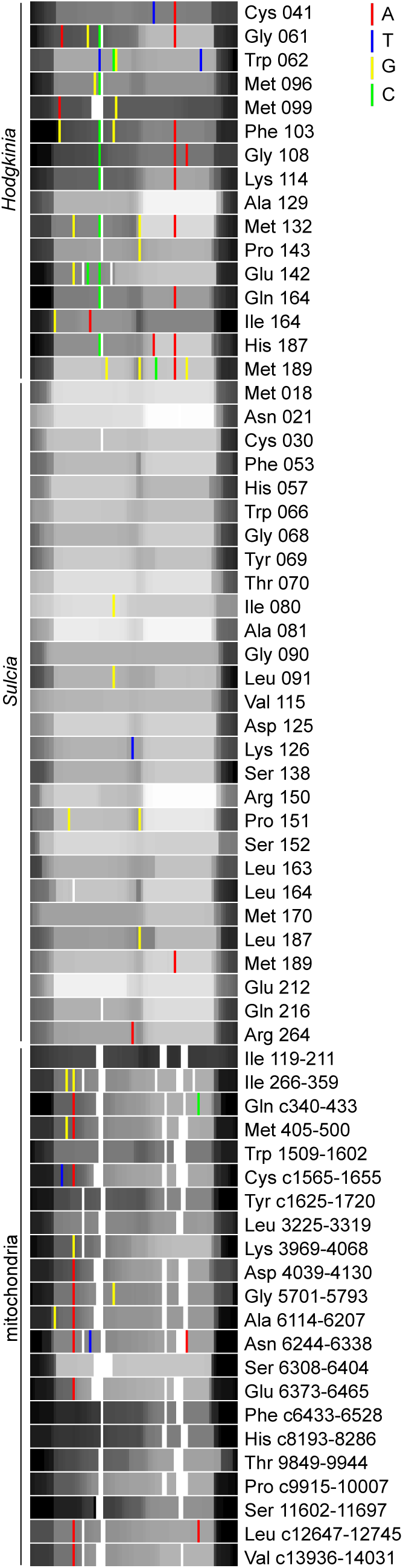
Expression level of individual tRNAs shown with polymorphic sites that have frequencies of greater than 2%. The per-base read depth was log transformed and is shown on a 0-255 color scale, making even large expression level differences difficult to distinguish by eye. Low expression is shown in black, high expression in white. Polymorphic sites are colored according to their genomic sequence. Ten bases of leader and trailer are shown as in Fig. 5. Gaps are shown in white and are apparent primarily in mtRNAs. See supplementary table S2 for gene name descriptions.

**Figure 6.**
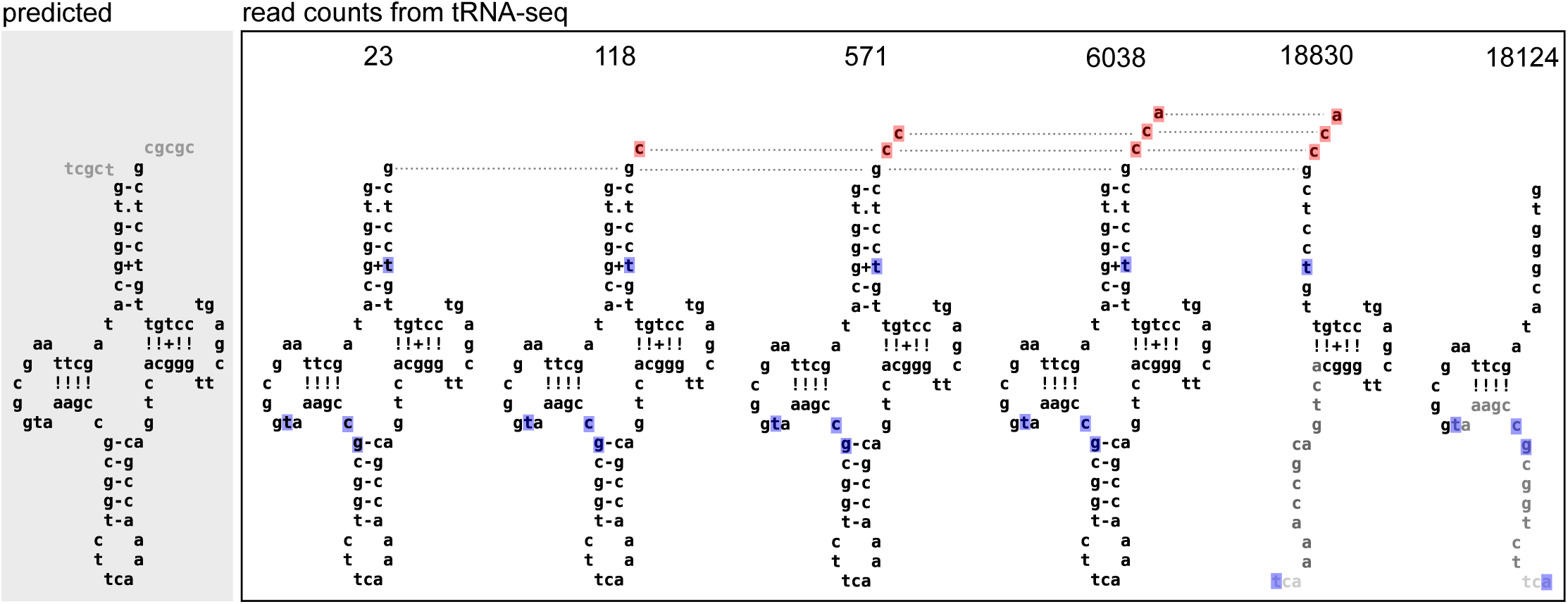
tRNA processing occurs in a stepwise manner, but full-length tRNAs comprise a small minority of the total reads. The majority of reads mapping to the *Hodgkinia* tRNA Trp062 gene (51,683) map to one of the secondary structures shown. Polymorphic sites (>2%) are shown in blue (RNA modifications) or red (CCA addition). tRNA halves are colored to indicate common sites of RNA degradation, where black letters indicate the highest read depth.

### Discovery of unannotated RNase P and tmRNA genes in *Hodgkinia*

By aligning small RNA reads to the *Hodgkinia* genome we found expression of previously unannotated RNase P (*rnpB*) and tmRNA (*ssrA*) genes at positions 25448-25794 and 92713-93140 in the *Hodgkinia* NC_012960.1 genome. Given that the 5’ end of *Hodgkinia* tRNAs are correctly processed (see the next section) and that we cannot find any other RNA nucleases in *Hodgkinia*, it seems likely that this RNase P is responsible for the observed tRNA processing. The permuted tmRNA is coded for in the reverse direction, on the anti-sense strand (Fig. S2). All components typically conserved in tmRNA structure can be found in the proposed tmRNA gene, however the peptide tag does not end in the typically conserved YALAA sequence (54, 55). The coding RNA and acceptor RNAs are separated by a 129nt intervening sequence containing complementary sequences needed for folding, and very few reads map to this region. Polymorphic sites indicate CCA addition at the 3’ end of both the coding and acceptor RNAs, further supporting the tmRNA’s likely functionality (56), especially given the presence of its protein ligand SmpB in at least two *Hodgkinia* genomes from different cicada species (57). We also observe reads of varying length at the ends of the tmRNA gene, indicating that end-trimming probably occurs.

### 5’ end processing of *Hodgkinia* and *Sulcia* tRNAs

Many reads aligning to *Hodgkinia* and *Sulcia* tRNA genes extend past the predicted gene boundary, suggesting that they are transcribed with 5’ leaders and that these extra nucleotides are trimmed off (Fig. 4). We previously predicted the presence of the RNA moiety of RNase P RNA in *Sulcia* (16), and we now predict the presence of this gene in *Hodgkinia* as described above. To help verify the activity of both the *Hodgkinia* and *Sulcia* RNase P RNAs, we created two pools of Illumina-compatible small RNA libraries. In the creation of both libraries, RNA adapter sequences are ligated directly to the small RNA pools at the 5’ and 3’ ends. Adapter ligation can be blocked by either a tri- or diphosphorylated 5’ RNA ends, but a functional RNase P will generate 5’ monophosphate ends which are active for ligation (Kazantsev and Pace 2006). By splitting one pool of small RNAs into two groups, one treated with Tobacco Acid Pyrophosphatase (TAP, which will generate 5’ monophosphates) and one untreated, we tested the 5’ processed state of bacteriome tRNAs (24, 30, 58). In both *Hodgkinia* and *Sulcia*, we found no difference between the tRNA sets from each library (Spearman’s rank correlation, p < 0.005; Table S5), suggesting that the 5’ ends of tRNAs are monophosphorylated in the cicada bacteriome. This is consistent with the presence of an active RNase P enzyme in both bacterial endosymbionts.

### The 3’ ends of *Hodgkinia* and *Sulcia* tRNAs are correctly processed

The processing of tRNA 3’ ends is more complicated than the processing of 5’ ends. If a 3’ CCA is not encoded on the genome (and the majority of *Sulcia* and *Hodgkinia* tRNA genes do not have encoded CCA ends), one must be added by a CCA transferase enzyme after the 3’ trailer sequence is processed off by various RNA nucleases. Consistent with the presence of 3’ trailer sequences, we find reads extending past the predicted 3’ boundary of *Hodgkinia* and *Sulcia* tRNAs (Figs. 4 and 6). *Sulcia* contains a putative ribonuclease (ACU52822.1) that could potentially process the 3’ trailer, although the gene is most similar to RNase Y, which is involved in mRNA decay (59). *Hodgkina* contains no such RNA nuclease candidates. We also observe reads ending in C, CC, and CCA that map to *Sulcia* and *Hodgkinia* tRNAs genes, indicating that each nucleotide of the terminal CCA is added one at a time to the 3’ end of transcripts lacking 3’ trailers (Fig. 6). *Sulcia* contains a tRNA CCA nucleotidyl transferase, but, again, *Hodgkinia* does not. Our mRNA-Seq data show upregulation of a tRNA CCA nucleotidyl transferase encoded on the cicada genome in the bacteriome tissue, although we do not know if this enzyme is active on *Hodgkinia* tRNAs. However, in plants, mammals, and yeast, isoforms of this protein are localized to both the cytoplasm and organelle (60–62), suggesting that this enzyme might be targeted to different cellular compartments in cicada cells.

### tRNA modification occurs in *Hodgkinia* and *Sulcia*

The *Hodgkinia* genome encodes only three genes known to be involved in tRNA modification, all of which are predicted to act on U34: *mnmA, mnmG* (*gidA*)*, and mnmE* (*trmE*) (16). MnmA catalyzes the 2-thiolation of U to s2U; MnmG and MnmE form a dimer that catalyzes the conversion of s2U to nm5s2U (31). The *Sulcia* genome encodes these three genes, along with *truA* and *tilS* (16). TruA modifies U38-U40 to pseudouridine and TilS converts C34 to I34, enabling the specific recognition of Met versus Ile anticodons (31). We find sequence polymorphisms—which we interpret as potential base modifications (63–66)—at several sites other than the expected position 34 in *Hodgkinia* (positions 1-4, 6, 7, 9, 15, 16, 18, 20, 23, 26, 27, 37, 43, 46, 49, 57, 58, 62, and 68) and the expected positions 34 and 38-40 in *Sulcia* (positions 7, 26, 37, and 58) (Fig. 5, Table S6). For a position to be called polymorphic, we required at least a 10X read depth and greater than 2% polymorphism at the modified site. Interestingly, *Hodgkinia* tRNAs are more highly modified than *Sulcia* tRNAs in both the diversity of modification and in the total number of tRNAs modified. Of *Hodgkinia’s* 16 tRNAs, 15 have at least one putatively modified site, versus 8 of 28 in *Sulcia* and 10 of 22 in the mitochondrial genome (Fig. 5, Table S6).

### Cicadas upregulate some aminoacyl tRNA synthetase transcripts in bacteriome tissues, but few HGT candidates

We sequenced cicada mRNAs from four insects in search of tRNA processing genes that could complement those missing from the *Hodgkinia* and *Sulcia* genomes. Sequencing eukaryotic RNA has proven to be a conservative and reliable method for finding functional HGT events in insect genomes (20, 45, 47). A total of 189,137 genes were assembled by Trinity from a combined 255,193,489 100bp reads. The longest and mean contig lengths were 18,931 bp and 765 bp, respectively. Transrate properly mapped 78% percent of the paired-end reads, giving an overall assembly score of 0.21. A total of 21,651 non-redundant annotated protein coding genes were found on the assembled contigs using Trinotate (supplementary material S1). BUSCO orthologs were searched and 75% of the core arthropod genes were found in our assembly, a typical value for hemipteran genomes (19). We used edgeR to identify 1,778 genes (supplementary material S2) that were differentially expressed between bacteriome and insect tissues (p ≤ 0.05). Of these 1,778 differentially expressed genes (DEGs), 1,211 had higher expression values in the bacteriome samples (supplementary material S3). Most DEGs were not annotated by Trinotate and remained hypothetical proteins. Only nine genes upregulated in the bacteriome were annotated with a function involving tRNA maturation or charging; five copies of D-tyrosyl-tRNA deacylase, two copies of aminoacyl tRNA synthase complex-interacting multifunctional protein, tRNA uracil(54)-C(5)-methyltransferase B, and tRNA modification GTPase MnmE (Table S7). The gene expression profiles were wildly different between insects (Fig. 7) so we examined the most differentially expressed genes (top quartile) which had non-zero CPM values for at least three of the four replicates and were not identified as differentially expressed by edgeR. Of the 205 tRNA maturation genes identified by the Trinotate annotation, nine genes not identified by edgeR were highly expressed in the bacteriome, but not so in other cicada tissues (Fig. 7). These genes are expressed 50-2,586 fold higher in bacteriomes than in other tissues. Some of the genes highly expressed in bacteriomes are complementary with the functions missing from the *Hodgkinia* and *Sulcia* genomes, including the arginine, cysteine, and serine aaRSs (Table 1).

**Figure 7.**
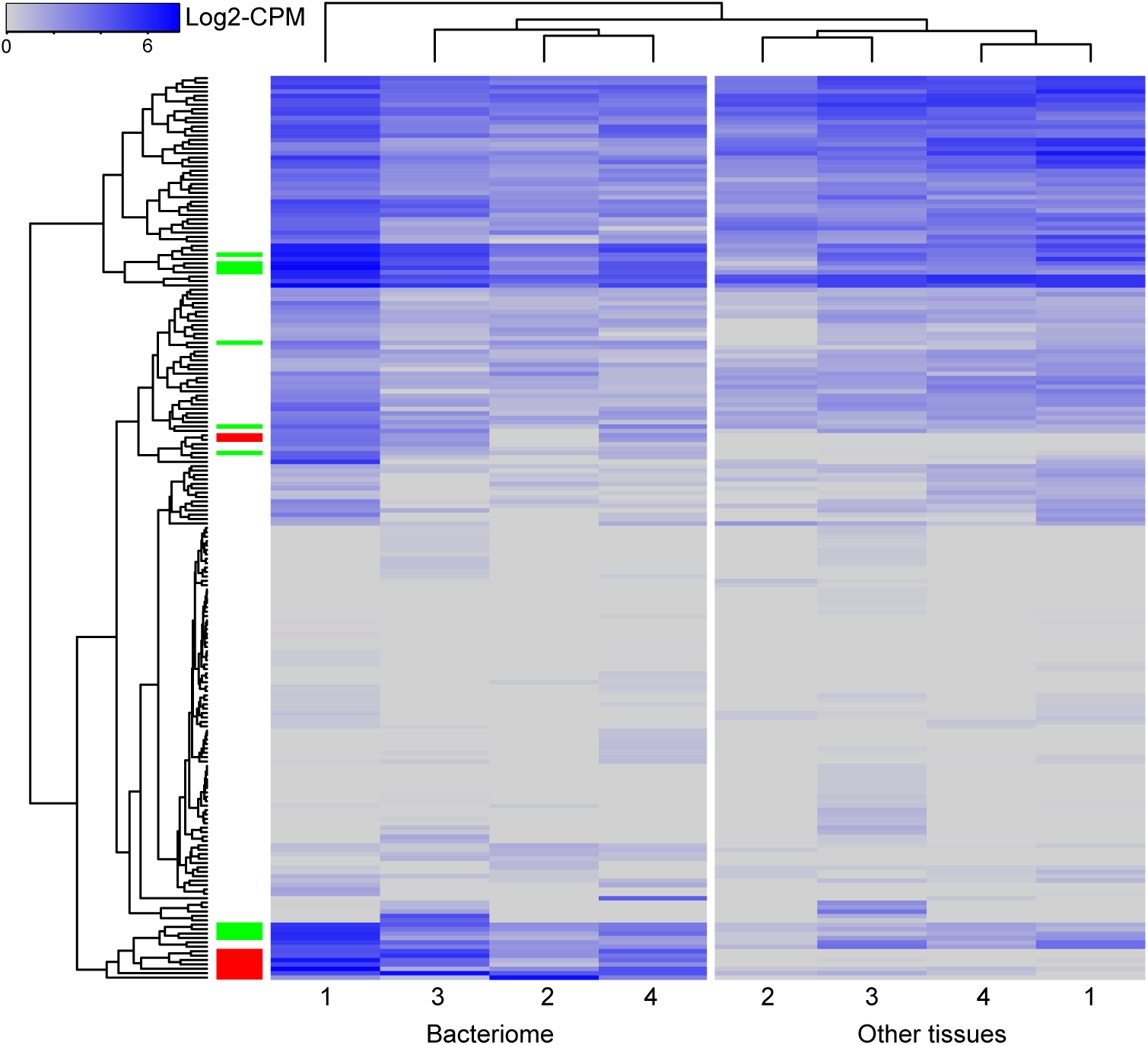
Host tRNA processing genes are rarely over-expressed in bacteriome tissues. Expression of Trinity assembled genes whose Trinotate annotations involve tRNA processing are shown for bacteriome and body tissues for replicate insects 1-4. Genes and samples are clustered by Euclidian distance in R. Differentially expressed genes (edgeR, p>0.05) are indicated by a red block in the left most column and genes differentially expressed, but not significantly so, are colored green.

**Table 1.**
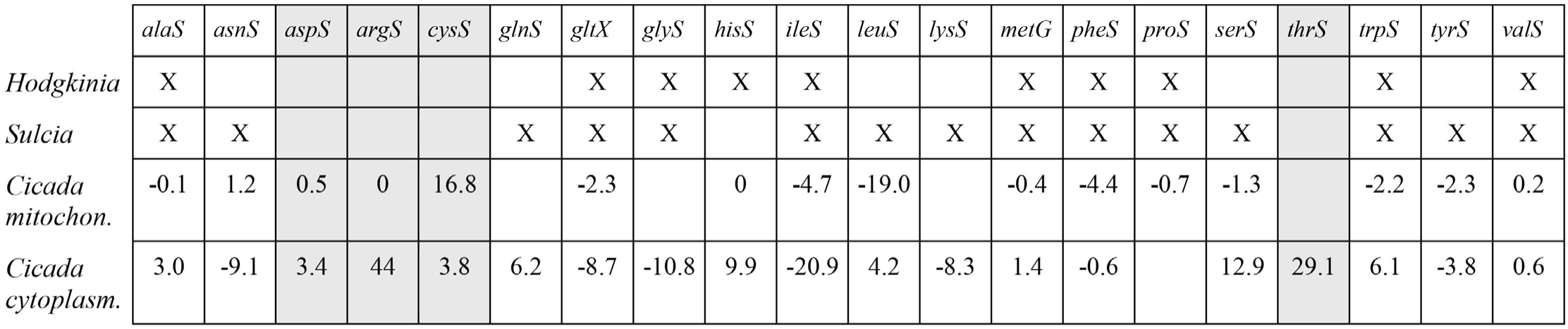
Expression of cicada aaRS genes in complementing *Sulcia* and *Hodgkinia.* The presence of aaRS genes in *Hodgkinia* and *Sulcia* genomes are indicated by an “X”. Differential CPM expression values of cicada aaRS genes are shown, where positive values indicate overexpression in the bacteriome (log2-CPM in bacteriome minus log2-CPM in other tissues).

Five cicada genes were identified as potential horizontally transferred genes (HTGs) from bacterial sources (Table S8), including pectin lyase (*pel*), two copies of a transcription regulatory gene (yebC), an AAA-ATPase, and ribosome recycling factor (frr). None of these genes are predicted to play direct roles in tRNA production or maturation. The functions of these are currently unknown. Overall, the cicada genome seems to contain few genes from HGT that obviously complement lost endosymbiont genes (with the possible exception of frr), in contrast to several related sap-feeding insects (20, 45, 47, 48).

### Methodological caveats and complexities

We found several unusual results while analyzing our data. First, we isolated highly abundant transcripts containing predicted tRNAs not belonging exclusively to *Hodgkinia*, *Sulcia*, or host tRNA genes. In these cases, each half of the transcript aligned to separate genomic locations of the endosymbionts, or even the genomes of separate organisms (half to *Sulcia* and half to *Hodgkinia*). In all cases, these were tRNA-like sequences that were joined near the anticodon. We could not amplify these RNAs from total RNA using gene-specific RT-PCR and thus concluded that they are a byproduct of the RNA ligation steps of the library preparation. This serves as a cautionary result of this method. Despite this, and in all cases, true *Hodgknina* and *Sulcia* tRNAs were also amplified, cloned, and sequenced as positive controls (see Table S3 for primer sequences). Second, we found tRNA modification patterns that do not correlate with the functional capabilities of *Hodgkinia* and *Sulcia*, and reasoned that modified nucleosides could disrupt reverse transcriptase during library preparation (66). These cDNAs will not contain both primer binding sites (adapters) and will not be amplified during the PCR step of the library preparation. Our protocol thereby selectively enriches for non-modified tRNAs. Since we find abundant tRNA sequences with polymorphism at conventionally modified sites, it seems likely that reverse-transcriptase can proceed over some modifications, consistent with previous findings (63–68). If this impacts our data significantly, we would predict that i) our tRNA abundance ranking may be incorrect and that tRNAs with extremely low coverage (i.e., *Hodgkinia* tRNAs) may have modifications besides those that we describe here (66) and thus were not amplified.

## DISCUSSION

Massive gene loss shapes the genomes of bacterial symbionts that adopt a long-term intracellular lifestyle. This process often results in metabolic dependencies between endosymbiont and its host. However, the loss of genes essential for transcription, translation, and replication is rare, occurring in only the most gene-poor bacterial genomes. How these organisms compensate for the loss of these genes is unknown. Here we show that despite seemingly incomplete symbiont capabilities to complete central cellular processes, *Hodgkinia* and *Sulcia* still produce mature tRNAs. We predict that completion of these processes is achieved through direct symbiont-host complementation.

### 5’ tRNA processing can be explained from the genomes of *Sulcia* and *Hodgkinia*, but 3’ processing cannot

We did not computationally identify RNase P in our first *Hodgkinia* genome annotation (16), but here we report the presence of this gene in the *Hodgkinia* genome. While still apparently lacking the protein component (*rnpA*), trimming of 5’ tRNA leaders could still occur with only the RNA component because it is a ribozyme (69). The 5’ processing of both *Hodgkinia* and *Sulcia* tRNAs can now be explained by normal cellular processes.

The way that *Sulcia* and *Hodgkinia* trim and process the 3’ ends of their tRNAs is less clear. Cleavage of the 3’ trailer from pre-tRNAs can be accomplished by a variety of redundant exo- and/or endonucleases (25), none of which are encoded in *Sulcia* or *Hodgkinia.* In *E. coli*, RNase PH, RNase T, RNase D and RNase II can all trim back the 3’ end of pre-tRNAs (25). In *Sulcia*, we identify a RNA nuclease with similarity to RNase Y. Although RNase Y is typically thought to initiate mRNA decay, it is also implicated in multiple RNA processing tasks (59). No such putative enzyme can be found in the *D. semicinta Hodgkinia* genome, although a gene for the endoribonuclease YbeY is present in *Hodgkinia* genomes from other cicada species (14). After 3’ tRNA trailers are trimmed, a terminal CCA is added by a CCA-adding enzyme (30). This gene is conserved across all domains of life (70), including *Sulcia*, but is apparently missing in *Hodgkinia.* We have identified transcripts belonging to *Hodgkinia* that have terminal C, CC, and CCAs. The presence of these variants indicates that *Hodgkinia* tRNAs are exposed to an active CCA-adding enzyme. One potential candidate is the host genome-encoded mitochondrial copy that is upregulated in bacteriome tissues (Table S7). Mitochondrial CCA-adding enzymes are known to have broad specificity and functionality (60, 71), and so may work on *Hodgkinia* tRNAs if directed to *Hodgkinia* cells by the host.

### Base modification, even if off-target and not biologically relevant, cannot be explained by the genes encoded in the *Sulcia* and *Hodgkinia* genomes

Base modifications are essential for tRNA aminoacylation and codon recognition, and have been well described in previous work (30), including the bacterial endosymbiont *Buchnera* (64). *Buchnera’s* tRNAs are post-transcriptionally modified at N37 and N34. Most of the observed modifications can be performed by enzymes encoded on the *Buchnera* genome (*miaA, miaB, rimN, trmD, tadA, queA, mnmA*, *mnmE, iscS, tilS*, and *gidA*), although an incomplete gene pathway is present for the Lys^TTT^-mnm^5^s^2^U observed in the tRNA sequencing data and several last-step enzymes are missing for N34 modifications. It is therefore a bit surprising that tRNA modifications are detected in *Hodgkinia* and *Sulcia* when the genes for these modification enzymes are not. On the other hand, edits at these sites are common in many species and would be expected to be observed in any normal bacteria with a reasonable genome. We note that *Hodgkinia* and mitochondrial tRNAs have more modifications in common than do *Hodgkinia* and *Sulcia* (Fig. 5, Table S6). Again, it seems possible that some of the host tRNA modification enzymes are active in *Hodgkinia* and *Sulcia*, although it is not clear if these modifications are biologically relevant.

### Translational machinery is shared between host and organelle in eukaryotes

The most gene-rich mitochondrial genomes of the Jakobid protists look very much like endosymbiont genomes, and contain a full-fledged set of about 30 tRNA genes (37). In contrast, the most gene-poor mitochondrial genomes of some trypanosomatides and alveolates contain no tRNA genes (72). The range is similar in plastids, from 1-30 tRNA genes (73, 74). The sets of retained tRNAs in *Hodgkinia* and *Tremblaya* overlap with organelles, but there are considerable differences (Table S1). Of 22 tRNA anticodon species in *Hodgkinia* and *Tremblaya* from the mealybug *P. citri*, only trnA^UGC^, trnI^GAU^, trnM^CAU^, and trnF^UGC^ are present in both genomes (Table S1). The tRNA gene conservation of *Hodgkinia*, mitochondria, and plastids is strikingly similar (75, 76).

Unlike in insect endosymbionts, organellar aaRSs have been completely transferred to the nuclear genome (77–79). The processes involved in aaRS and tRNA import into organelles are complex and are reviewed elsewhere (77, 80, 81). The mechanisms for localizing charged tRNAs to the organelle are diverse and organism specific. In humans, for example, all aaRSs except GlnRs and LysRS are bacterially-derived and targeted specifically to mitochondria to charge mitochondrially-encoded tRNAs (82). In contrast, only 45 aaRS genes are expressed from the *A. thaliana* genome; 21 are found only in the cytoplasm, 21 are dually-targeted, 2 are chloroplast specific, and 1 is targeted to all three cellular compartments (65). The mitochondrial genomes of apicomplexans are missing tRNAs and aaRSs. They must import aminoacylated tRNAs that were charged in the cytoplasm (83). It is worth noting that the mitochondrial and plastid genomes of *A. thaliana* contain 22 and 30 tRNA genes, yet cytosolic tRNAs are still imported (84). The import of seemingly unnecessary tRNAs occurs quite frequently, and in most cases, the role of redundant tRNAs in organelles is unknown (81). Our results strongly suggest that *Hodgkinia* and possibly *Sulcia* import tRNAs from their host cicada.

Despite the ancient nature and massive genetic integration of organelle with host (85, 86), most mitochondria and plastids retain genomes, and thus some level of genetic autonomy. Gene retention patterns in the genomes of highly reduced bacterial symbionts also suggest a hurdle to giving up independence of some processes to the host, especially transcription, translation, and replication (11, 12). Although the *Hodgkinia* genome is lacking many genes involved in these processes, our results here indicate that it is likely complemented by its host to perform them. While these results support the idea that some obligate symbioses may undergo major transitions to become a highly integrated unit (87), other recent data from other cicadas show that these integrated units are not inevitably stable—sometimes *Hodgkinia* is lost and replaced by a new fungal symbiont (88).

## MATERIALS AND METHODS

### Sequencing small RNAs

The bacteriomes of three wild caught female *Diceroprocta semicincta* collected around Tucson, Arizona in July, 2010 and July, 2012 were dissected and stored in RNA-Later (Ambion). Total RNA was later purified using the Roche High Pure miRNA Isolation kit following the total RNA protocol. Small RNAs were isolated with the same kit, but following the 2-column protocol for <100nt RNAs. RNA adapters were ligated to the 5’ and 3’ ends of the small RNAs using the Scriptminer™ Small RNA-Seq Library Preparation Kit from Epicenter. One index was treated with the supplied TAP enzyme to reduce the 5’ end to a monophosphate. Reverse transcription was done with an adapter specific primer and each library was subjected to 15 rounds of PCR using FailSafe PCR Enzyme Mix (Epicenter) and the supplied primers (94°C for 15 sec, 55°C for 5 sec, 65°C for 10 sec). PCR bands of approximate size 50-300nt (including 113nt adapters) were cut from an 8% polyacrylamide gel after staining with SYBR^®^ Safe (Invitrogen), and visualized on a standard UV transilluminator. The gel was shredded using a 0.5 mL tube with needle holes in the bottom, and eluted with 300 uL 0.5 M ammonium acetate for 3.5 hours at 37°C. The liquid was separated from gel particles using a 0.22 micron sterile filter and DNA was purified by standard isopropanol precipitation. Bioanalyzer traces of both libraries show DNA of about 100-275bp at sufficient concentration for Illumina sequencing. 226,712,931 100nt single-end reads were generated on three HiSeq lanes at the UC Berkeley Vincent J. Coates Genomics Sequencing Laboratory.

### Read processing for small RNAseq

Adapter sequences were trimmed using Cutadapt version 1.0 with options -a AGAT CGGAAGAGCACACGTCTGAACTCCAGTCAC -g AATGATACGGCGACCACCGACAGGT TCAGAGTTCTACAGTCCGACGATC -O 7 (89). Then, reads less than 18nt in length were discarded. Reads were quality filtered using FASTX-Toolkit version 0.0.12 (http://hannonlab.cshl.edu/fastx_toolkit/) so that reads with a quality score less than 20 over more than 10% of the read were discarded (fastq_quality_filter -q 20 -p 90). Datasets with reads of length 18-90, 48-90, and 70-100nt were generated using a custom Perl script. The size of 18nt was chosen because the identical matches up to 16nt in length can be found between different symbiont tRNA genes. The size 48nt was chosen because the shortest tRNAs are about that length (90). At this point, each of these datasets were used for mapping to *Hodgkinia* and *Sulcia* genomes and tRNA genes using either bowtie-1.0.0, with settings –best –maqerr 150 –seedlen 18 or bwa-0.7.5 aln, with settings -n 0.08 -i 2 (91, 92).

### De novo small RNA discovery

Identical reads from the 48-90nt dataset were compressed using FASTX-Toolkit (fastx-collapser). The majority of collapsed sequences were comprised of only one read, so a cutoff value was determined arbitrarily using a histogram of sequence coverage. The distribution of sequence coverage between 100X and 2E6X was quite even. The number of sequences with coverage from 100X-1X increases dramatically, so that there were 15,115 collapsed sequences with coverage higher than 100X and 6,478,420 collapsed sequences with coverage less than 100X. Therefore, all sequences comprised of less than 100 identical reads were discarded (6,463,305 sequences). The remaining 15,115 sequences were split into two sets: reads with BLASTN hits to *Hodgkinia* and *Sulcia* tRNA genes, and reads without hits (blastall 2.2.25, blastn -e 1E-25). Sequences that did not align with known, endosymbiont tRNAs were then aligned to the *Hodgkinia* and *Sulcia* full genome sequences (blastn -e 1E-10). The remaining sequences that did not align to the bacterial genomes were considered cicada sequences, and tRNAs were predicted using tRNAscan-SE 1.21 and ARAGORN 1.2.34 (93, 94). Nearly identical sequences were grouped into contigs using CAP3 (95). Collapsed sequences with different anticodons, 5’ leaders or 3’ trailers that assembled together in CAP3 were separated into their own contigs for bowtie-0.12.7 and BWA-0.5.9 alignments using custom Perl scripts.

### Comparing TAP treated to untreated libraries

Differential expression between libraries was compared using by expression rank changes and edgeR differential expression analysis. Reads from the 20-100nt and 70-100nt datasets were mapped to a multi-fasta file containing *Hodgkinia, Sulcia*, and mitochondrial tRNA genes plus 15bp of genome sequence flanking the gene using bowtie-0.12.7 with the -f, -S, and -n 3 options. tRNA abundance rankings were generated from the *.sam files by simply counting the number of reads that mapped to each tRNA sequence listed in Fig. 3. The order of tRNA coverage was compared between indexes using Spearman-rank correlation (Table S5). Trinity v20140717 packages align_and_estimate_ abundance.pl, abundance_estimates_to_matrix.pl, run_DE_ analysis.pl, and analyze_diff_expr.pl scripts were used to compare differential transcription with parameters (-SS_lib_type F –est_method RSEM –aln_method bowtie –seedlen 18 –maqerr 150 –best). Using the de novo approach separately for library 1 and 2, with 48-100nt reads, we normalized tRNA coverage (number of reads per tRNA/total number of reads mapping to all tRNAs). A ratio of difference between library 1 and 2 coverage was calculated for each tRNA. For all values less than zero, the inverse was taken and multiplied by −1. In this way, we tried to capture the relative difference in expression for all tRNAs from all organisms. These data were tabulated so that source organism, paired amino acid type, anticodon sequence, and relative expression change for every tRNA were in one row. In R lm() and anova() were used to model factor effects on tRNA expression (Tables S5 and S9).

### mRNA-seq analyses

Illumina reads from cicada bacteriome and non-bacteriome tissues (SRR952383) were pooled with reads from an additional three wild caught female *Diceroprocta semicincta* collected around Tucson, Arizona in July, 2012 and assembled using Trinity v2.1.1 using kmer_ length = 25 and min_contig_length = 48 (96). The edgeR package was used to analyze differential expression with RSEM quantification and bowtie alignments (97). The quality of the assembly was assessed for quality with Transrate v1.0.2 and completeness with BUSCO v1.22. Assembled transcripts matching *Sulcia* and *Hodgkinia* with 100% identity were removed by mapping with bwa-mem v07.12. The remaining transcripts were annotated using Trinotate v2.0.2. Potential horizontally transferred genes (HTGs) were identified from the cicada transcripts with BlastP searches against the bacterial nr database and verified against known HTGs from the leafhopper *Macrosteles quadrilineatus* (Mao & Bennett unpub.) since none currently exists for *D. semicincta.* Of the 398,377 Trinotate annotated gene isoforms, 115,253 have blast hits in the SwissProt, TrEMBL, or EggNOG databases. These 114,556 isoforms are located on the 21,651 transcripts reported in supplementary material S1. 1,284 contaminated transcripts were removed from the Trinity assembly and Trinotate annotation list following identification during submission to NCBI’s transcript sequence archive (TSA). Genes involved in tRNA processing were identified by searching for “tRNA” or “transfer RNA” in the Trinotate annotation. Expression values were analyzed and visualized in R using the edgeR, trinotateR, and ggplots2 packages.

### Cloning and sequencing of prepared libraries and tRNAs

Cloning was done using Invitrogen’s TOPO TA cloning kit with OneShot TOP10 chemically competent cells using standard procedures. Primers designed to be specific to the tRNA of interest were used to prime reverse transcription using Invitrogen’s superscript III First-Strand Synthesis kit. NEB OneTaq was used in end-point PCR prior to cloning (standard reaction with 2uL RT product and 40 cycles). Promega PCR ladder and NEB 6X loading dye was used to visualize PCR products prior to cloning. Plasmids were purified using Omega’s Plasmid Mini Kit and sequencing was done with the standard M13F primer.

### Bioinformatics

Complete bacterial genome sequences were downloaded from ftp://ftp.ncbi.nlm.nih.gov/genomes/Bacteria/all.fna.tar.gz. Chromosomal sequences were searched for tRNA genes using tRNAscan-SE 1.21 using the bacterial model (94). Genomic GC contents and 4-box family tRNA gene counts were calculated with custom PERL scripts. 6-box families were included in the analysis. tRNA redundancy is simply calculated by dividing the number of 4-box family tRNA genes by the number of 4-box families.

## ACKNOWLEDGMENTS

The authors thank the McCutcheon lab members and Juan Alfanzo for technical assistance and conceptual feedback, Todd Lowe and Patricia Chan for running a custom Infernal search on the *Hodgkinia* genome, Filip Husník for contributions to Table S1, and Bodil Cass for cicada collection. This work was supported by National Science Foundation (NSF) grants IOS-1256680 and IOS-1553529, and by the National Aeronautics and Space Administration Astrobiology Institute Award NNA15BB04A. Raw Illumina reads are available in NCBI’s SRA database for the small RNA (SRR5081152-5081157) and transcriptome (SRS470226 and SRR5060328-5060333) datasets. Assembled transcript sequences are available from the NCBI TSA database (GGPH00000000.1).

## SUPPLEMENTARY MATERIAL

**Supplementary material S1.** Trinotate annotations for all transcripts assembled using Trinity. Available at https://doi.org/10.6084/m9.figshare.6687089.v1.

**Supplementary material S2.** Expression level of gene-level transcripts that were differentially expressed between bacteriome and other cicada tissues. Shown as log2 fold-change. Available at https://doi.org/10.6084/m9.figshare.6687089.v1.

**Supplementary material S3.** Trinotate annotations for transcripts there are expressed higher in cicada bacteriome tissues than in other cicada tissues. Available at https://doi.org/10.6084/m9.figshare.6687089.v1.

